# SENSITIVITY OF MEDIAL/LATERAL BALANCE CONTROL TO VISUAL DISTURBANCES WHILE WALKING IN YOUNG AND OLDER ADULTS

**DOI:** 10.1101/2024.12.10.627754

**Authors:** Stephen J. DiBianca, Hendrik Reimann, Julia Gray, Robert J. Peterka, John J. Jeka

**Affiliations:** University of Delaware, Kinesiology and Applied Physiology, Newark, DE, United States; Oregon Health & Science University, Portland, OR, United States

## Abstract

Humans integrate multiple sources of sensory information to estimate body orientation in space. Balance control experiments while standing provide evidence that the contributions of these sensory channels change under different conditions in a process called sensory reweighting. This study aims to address whether there is evidence for sensory reweighting while walking and explores age-related differences in medial/lateral balance control under visually perturbing walking conditions. Thirty young adults (18–35 years) and thirty older adults (55–79 years) walked on a self-paced treadmill within a virtual environment that delivered frontal plane multi-sine visual disturbances at three amplitudes (6°, 10°, and 15°). Frequency response functions were used to quantify visual sensitivity to balance disturbances, while spatiotemporal gait parameters (e.g., step width, step-width variability) were measured to assess balance control. Visual sensitivity decreased in both populations with increasing stimulus amplitude, analogous to the sensory reweighting hypothesis in balance control while standing. Despite the decrease in visual sensitivity, the compensatory upweighting of other sensory systems was not observed through measurements of remnant sway. Older adults exhibited higher visual sensitivity at all amplitudes compared to young adults, indicating a more sensitive response to visual disturbances to balance control. Both groups showed increases in step width and step width variability with higher visual amplitudes, with older adults demonstrating more pronounced effects. Weak correlations existed between changes in visual sensitivity and changes in step width and step width variability suggesting a limited interaction between sensory reweighting and gait stability.

## Introduction

Upright balance control is a continuous sensorimotor process integrating information from multiple sensory systems and coordinating motor outputs to maintain upright posture (Shanbhag et al., 2023). Sensory pathways from the visual, vestibular, and proprioceptive systems are integrated by the nervous system to provide an estimate of the body’s position and velocity in space (Mahboobin et al., 2009; Horak and MacPherson, 1996). The nervous system can be thought of as a maximum-likelihood integrator which takes a weighted combination of these position and velocity estimates from each sensory pathway to produce an optimal estimate that has lower variance than either of the individual sensory signals (Ernst and Banks, 2002). The optimized position and velocity sense informs the selection of motor outputs to maintain the center of mass over the base the support. This process of changing reliance to a sensory system to facilitate the estimation of the body’s motion state in space (i.e., position and velocity) is called sensory reweighting.

Experiments have extensively investigated sensory reweighting in standing balance control through a linear systems theory approach in which a participant is driven by a sensory stimulus and resulting body motion is measured. A frequency response function characterizes how the 1) sensitivity (gain) and 2) timing (phase) of the body motion change as a function of stimulus frequency. Probing all three sensory modalities either simultaneously (Hwang et al., 2014) or separately (Goodworth and Peterka, 2012; Jeka et al., 2010; Cenciarini and Peterka, 2006), this body of work has revealed that the nervous system systematically changes sensitivity (gain) of individual sensory channels to minimize variance that would drive excessive sway resulting in a loss of balance characterized by a stepping response. Optimal estimation and control theory predicts that an improved orientation estimate can be obtained by down-weighting the sensory system perturbed (van der Kooij et al., 2001) and up-weighting another to insure there is sufficient torque generated by the muscles to resist gravity (Cenciarini and Peterka, 2006 Oie et al., 2002). Computational findings from these principals indicate that the reweighting process that adjusts for an external perturbation is characterized by an increase in remnant body sway (van der Kooij et al., 2001). Remnant body sway is a measure of the magnitude of sway not correlated with the driving frequency of the perturbing sensory signal. These computational findings indicate that sensory noise provides the major contribution to remnant body sway in standing, such that as the sensitivity to a sensory modality decreases, the remnant body sway increases indicating an up-weighing of alternative sensory sources of orientation information (van der Kooij and Peterka, 2011).

Our first research question addresses whether there is evidence for sensory reweighting as a neural mechanism for balance control while walking. However, probing postural control during gait is not straightforward. Linear system methods have been fruitful in probing sensorimotor integration in standing since standing can be approximated as a linear stochastic system in which the base of support is fixed (Kiemel et al., 2002; Peterka, 2002). In walking, however, the base of support moves with each step and includes a nonlinear limit cycle where vestibular (Bent et al., 2004, Reimann et al., 2019), somatosensory (Duysens et al., 1995) and visual disturbances (Logan et al., 2010) affect body mechanics differently depending on the body’s configuration during stimulation (i.e., single vs double stance). For example, vestibular stimuli delivered at heel-strike compared to mid-stance or toe-off produce similar onset timing of muscle activation but different body responses depending on which balance mechanisms are available and for how long (Reimann et al., 2019). A stimulus at toe-off, for example, will result in a larger foot placement response on the next step compared to the same stimulus delivered right before heel strike. With plenty of time post stimulus to swing the leg outward, a significant lateral foot placement shift occurs with the toe-off stimulus. On the other hand, when the same vestibular stimulus is delivered right before heel strike, there is not sufficient time to shift the swing leg, which leads to a larger ankle response of the stance leg to produce more ankle torque to quickly adjust the center of pressure. It is currently unclear whether the nervous system adjusts sensitivity to a sensory stimulus during locomotion or maintains sensitivity and solely exploits the redundancy in degrees of freedom available for different motor strategies.

Our second research question explores if any differences exist in sensitivity to visual perturbations between healthy young adults versus older adults. Globally, 26.5% of older adults fall each year (Salari et al., 2022). The vestibular, proprioceptive, and visual system all degrade with age. Hair cell counts decline in the vestibular system (Merchant et al., 2000), leading to increases in vestibular motion detection thresholds after age 40 for yaw and roll rotations as well as horizontal and vertical translations (Bermúdez Rey et al., 2016). The sensitivity, acuity, and integration of the proprioceptive signal of the lower limbs for balance control declines with age (Henry and Baudry, 2019). Anatomical changes of the eyes occur in which the lens becomes thicker and loses elasticity, affecting the ability to focus on near and far-sighted objects (Saftori and Kwon, 2018). Furthermore, increases in visual motion detection thresholds have been observed in older adults (Conlon et al., 2017; Snowden and Kavanagh, 2006), although there is evidence that older adults can discriminate visual motion of high contrast better than young adults (Tadin and Blake, 2005). For standing balance, the process of visual reweighting has been shown to take longer in healthy and fall-prone older adults compared to young adults (Jeka et al., 2010) which can hinder the ability to recover from sudden environmental disturbances. We are currently unaware of what difference may exist in sensitivity to visual disturbances while walking. We hypothesize that older adults will be less able to limit visually-evoked disturbances to balance as the stimulus amplitude increases compared to younger adults.

As an exploratory analysis, we were interested in comparing sensitivity measures to spatiotemporal gait parameters. Walking stability has gained more attention in recent years since falls most commonly occur while walking (Kelsey et al., 2012). By categorizing changes in visual sensitivity as a proxy for sensory reweighting capabilities, we were interested in exploring if changes in sensitivity relate to stability. By adding a visual disturbance, we expected to destabilize participants. Medial/lateral (M/L) visual disturbances have been shown to increase average step width in both young and older adults (Kazanski et al., 2020, McAndrew et al., 2010). Furthermore, we expected to observe an increase in gait variability as an indicator of destabilization (Bruijn et al., 2013; Hausdorff, 2005; McAndrew et al., 2010). During unperturbed walking, older adults typically exhibit higher foot placement variability compared to young adults (Owings and Grabiner, 2004) and are at a higher risk of falling. Our motivation was to explore whether the degree of visual down-weighting had any predictive value on the changes in resulting step width and step placement variability. While measures of gait variability provide insight into the internal noise of the controller, variability alone does not imply instability (Bruijn et al., 2013). Since there is redundancy in the degrees of freedom available to motor strategies for upright balance control during walking, gait variability may indicate a healthy controller that can adjust and adapt to different environments, however too little or too much is associated with fall risk (Brach et al., 2005).

Because the neural control for walking mainly requires corrective action in the M/L direction, while passive dynamic mechanisms contribute to stabilization in the anterior/posterior direction (O’Connor and Kuo, 2009), we focused our gait parameters to those in M/L direction.

These M/L corrective actions, or balance mechanisms, include lateral foot placement of the swing leg towards a perceived fall (Bauby and Kuo, 2000; Wang and Srinivasan, 2014), lateral ankle modulation to generate torque under the stance foot (Reimann et al., 2017, Hof and Duysens, 2018), and a push-off of the trailing limb onto the stance limb (Reimann et al., 2018). The locomotion literature has largely focused on changes in foot placement in balance control during walking (Kazanski et al., 2020; Owings and Grabiner, 2004; Bruijn and van Dieën, 2018). Here, we compared whether visual sensitivity changes with stimulus amplitude affected the way these populations regulate step placement as characterized by average step width and step-width variability.

In this paper we probe the balance control system using a multi-sine visual stimulus that included a broad frequency range (0.025-0.475Hz) relevant to balance control during treadmill walking trials (>300 steps) in healthy young and older adults. The multi-sine stimulus tilted the visual scene in the medial/lateral direction at three different peak-to-peak amplitudes: 6, 10, and 15 degrees. We hypothesize that the sensitivity to the visual stimulus will decrease as the stimulus amplitude increases for both groups with the potential that the sensitivity changes may be indicative of sensory reweighting that reduces reliance on vision while increasing reliance on proprioceptive and vestibular information. We expect a decrease in visual sensitivity to be larger in young than in older adults given that young adults are expected to have more reliable proprioceptive and vestibular sensory information. We also hypothesize that remnant body sway will be larger in older than younger adults and will increase in both populations with increasing visual stimulus amplitude as the overall combination of sensory contributions to balance become less reliable.

## Research Design and Methods

Thirty healthy young participants (age range 18-35 years) and 30 healthy older participants (age range 55-79 years) participated in this study. Participant characteristics are shown in Table 1. A screening procedure excluded potential participants who: had head, neck, face, or lower extremity injury within 6 months prior to experiment date, were taking any medication that could affect balance, had any neurological or cardiovascular conditions, did not have normal or corrected to normal vision, were pregnant, and/or had a BMI over 30. Verbal and written consent were obtained from each participant. The experimental protocol was approved by the University of Delaware Institutional Review Board.

**Table 1:** Subject Characteristics. Data are presented as Mean ± Standard Deviation.

| Subject Characteristics (n = 60) | Healthy Young (n = 30) | Older Adults (n = 30) |
| --- | --- | --- |
| Age (years) | 26.7 ± 3.9 | 65.2 ± 5.1 |
| Body Mass (kg) | 72.0 ± 12.8 | 73.0 ± 14.4 |
| Height (m) | 1.7 ± 0.1 | 1.7 ± 0.1 |
| Sex (female/male) | 14/16 | 16/14 |

## Experimental Protocol & Setup

Figure 1 represents the experimental design. Participants first performed a set of psychophysical tests counterbalanced with the experimental design on the same day. The psychophysical tests, to be published separately, assessed the ability of participants to determine the direction of visual display movement during standing and walking. Participants walked on a self-paced, tied-belt treadmill (Bertec, Inc, Columbus, OH) inside a virtual environment displayed on a large dome that occupied the subjects’ full visual field (Figure 2). All participants performed an initial 15-minute familiarization block by walking on the self-paced treadmill while viewing a 15-degree pseudo-random visual perturbation which we refer to as the ‘visual multi-sine stimulus’ (defined below - Visual Multi-sine Stimulus). After familiarization, the experiment included six trials of walking while following a 0.8 Hz metronome with two beats per gait cycle, or 96 bpm, while viewing the multi-sine stimulus at 6, 10, and 15 degrees peak-to-peak amplitudes presented in a randomized order. Participants performed two trials at each stimulus amplitude. Each trial lasted 290 s and included 17 s of unperturbed walking followed by six 40-s duration cycles of the visual multi-sine stimulus.

**Figure 1:**
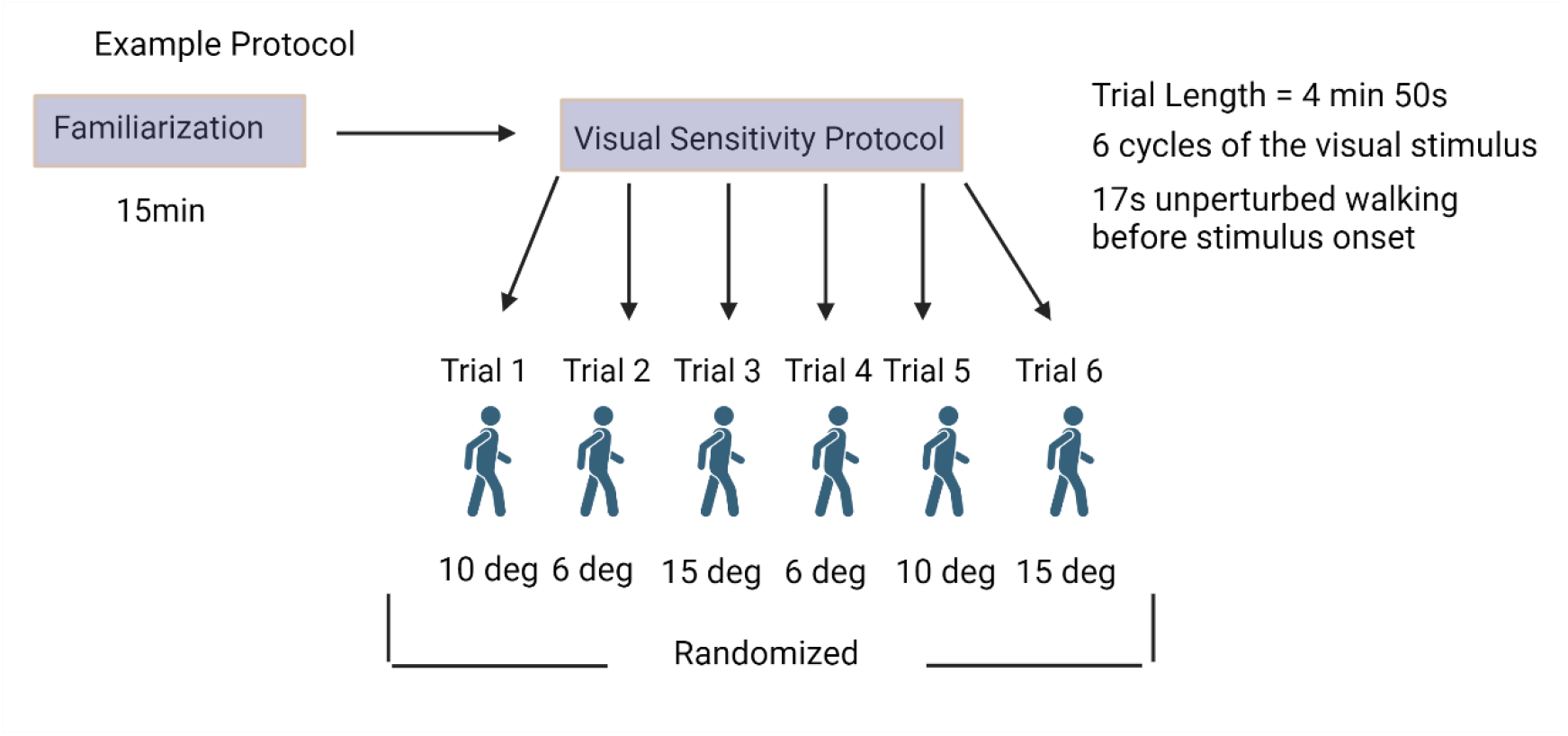
Shows the experimental design. Participants first experienced a familiarization period where they: 1) walked on the self-paced treadmill in the virtual environment, 2) were exposed to the visual multi-sine stimulus, and 3) assimilated to walking with the metronome cadence. After familiarization, participants experienced 6 trials of walking while viewing the visual multi-sine with a metronome, each 4 min 50 s long. The trials were randomized and included 2 trials of each stimulus amplitude. Created with BioRender.com

**Figure 2:**
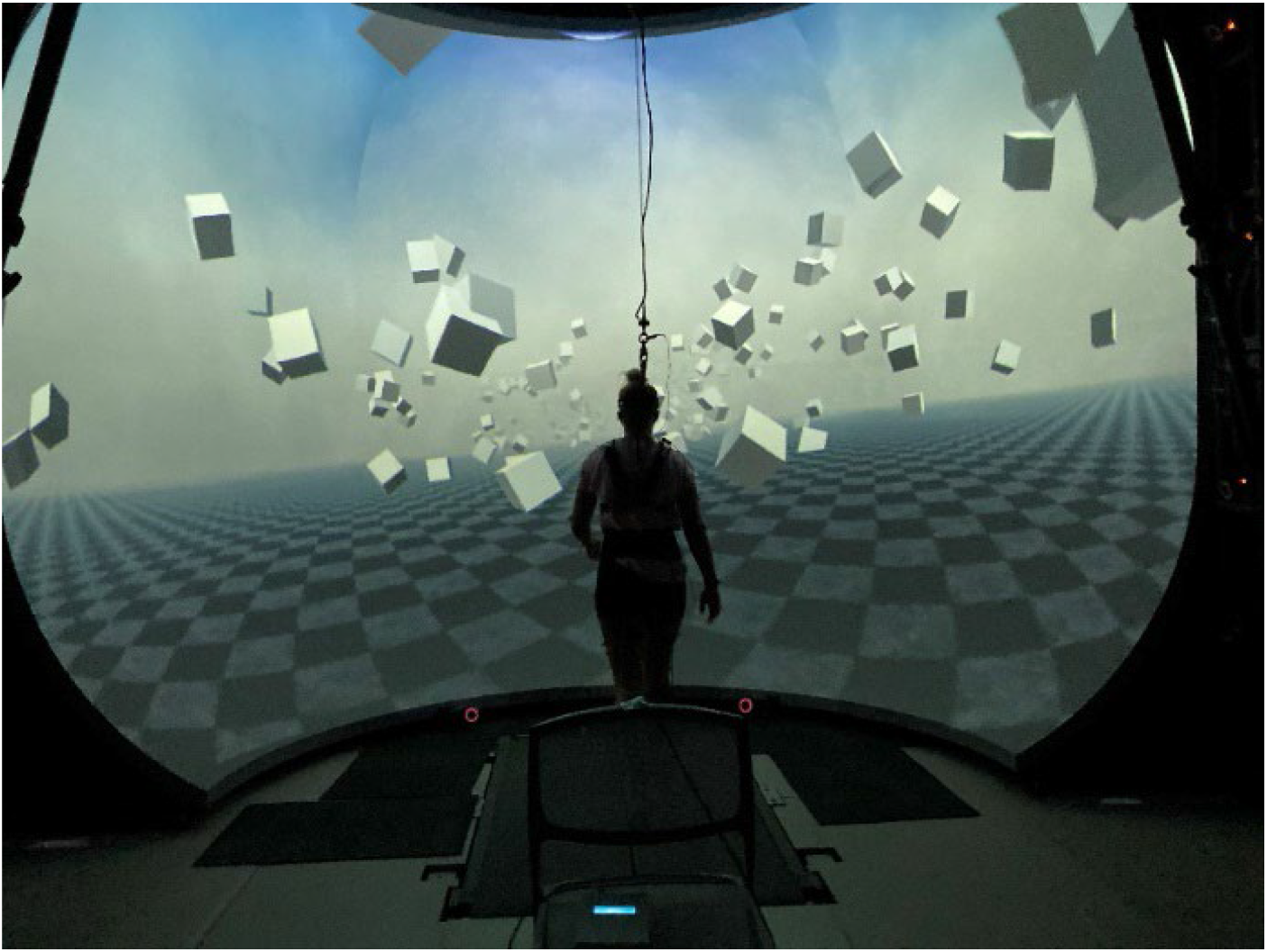
Displays the experimental set up in which healthy young and older adults walked on a self-paced treadmill in the virtual dome with the 44 reflective markers placed along the body. The visual scene tilted using the multi-sine stimulus.

Using the Plug-in Gait marker set by Davis et al. (1991), forty-four reflective markers were placed bilaterally on the subjects’ feet, shank, upper legs, hips, torso, upper arms, forearms, and hands, with six additional markers placed on the anterior thigh, anterior shank, and 5^th^ metatarsal. Marker coordinate positions were recorded using a Qualisys Motion Tracker System with 13 cameras at a sampling rate of 200 Hz. The self-paced control of the treadmill speed was implemented to maintain the subject in the center of the treadmill during walking trials. The control used a nonlinear proportional-integral-derivative (PID) controller algorithm implemented in Labview (National instruments Inc., Austin, TX) to keep the midpoint between the two reflective markers placed on the posterior iliac spine at the midline of the treadmill. Force plates integrated in the treadmill measured ground reaction forces and moments at a sampling rate of 1,000 Hz from both belts. Subjects wore a safety harness in the event of a fall, although none occurred.

## The Virtual Scene

Figure 2 shows a depiction of the virtual environment which was designed and implemented in Unity3d (Unity Technologies, San Francisco, CA). The virtual environment consisted of a tiled marble floor with floating cubes randomly distributed in a volume 0–10 m above the floor, located 2–17 m to each side of the midline, and infinitely into the distance forming a 4 m wide corridor for the subjects to walk through. A fog was displayed in the distance to obscure the limits of the tunnel and create a perception of an infinite walking path. The anterior/posterior movement of the virtual scene matched the speed of the treadmill with perspective in the virtual world linked to the midpoint between the two markers on the subject’s temples. This perspective was linked to the center of the floor horizon and was the axis of rotation for the visual multi-sine stimulus.

## Visual Multi-sine Stimulus

The visual multi-sine stimulus consisted of a combination of ten sinusoids at frequencies between 0.025 and 0.475 Hz (placed at odd harmonics of the 0.025 Hz fundamental), containing power at a wide range of frequencies typically observed in upright balance control (Yamamoto 2015). The individual sinusoids were phase-shifted, scaled and superposed to create a smooth multi-sine stimulus (Ouderaa et al., 1988) that gave a continuous seesaw like, side-to-side tilting motion of the virtual environment around an anterior/posterior axis at floor level. Figure 3A shows one 40-s cycle of the visual multi-sine with 15-degrees peak-to-peak amplitude and the corresponding amplitude spectrum (Figure 3B) with the 10 frequency components of the stimulus highlighted in red.

**Figure 3:**
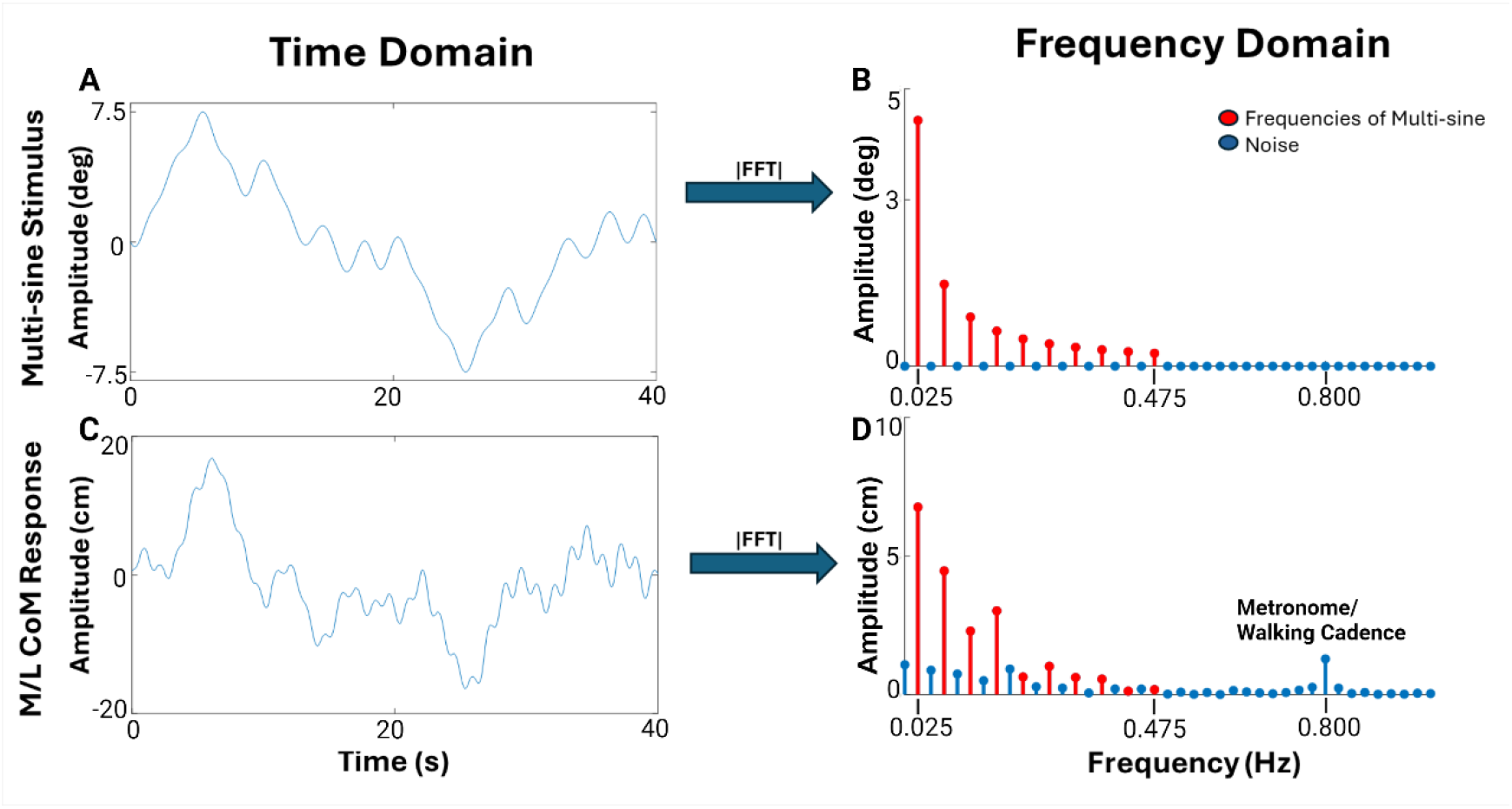
(A) shows one cycle of the multi-sine visual stimulus in the time domain. The x- axis represents time (s) and the y-axis represents amplitude (degrees). (B) shows the frequency domain amplitude spectrum of the multi-sine stimulus. Stimulus amplitude of the sinusoidal components can be seen at the 10 frequencies highlighted in red. These frequencies are odd harmonics of the 40-s stimulus cycle, spanning 1/40 Hz to 19/40 Hz. (C) shows a participant’s cycle-average M/L CoM response to walking while viewing the visual multi-sine in the time domain. (D) shows the amplitude spectrum of the CoM response. The highlighted red lines represent M/L sway amplitude (in cm) at the 10 driving frequencies of the visual multi-sine stimulus. Also evident is the amplitude of the M/L sway at the participants stepping frequency driven by the metronome at 0.8 Hz.

## Measuring Spatiotemporal Gait Parameters

Center of mass (CoM) position data was calculated based on a geometric model with 15 segments (pelvis, torso, head, thighs, lower legs, feet, upper arms, forearms, hands) and 38 degrees of freedom (DoF) in OpenSim (Delp et al., 2007; Seth et al., 2018) using an existing base model (Zajac et al., 1990). CoM movement was measured as the motion of the estimated CoM position in the M/L direction. CoM response was measured as the filtered CoM movement using a 6^th^ order low pass Butterworth filter with a cutoff frequency of 0.5 Hz. The CoM response assesses frequencies within the range of the visual multi-sine while filtering out the motion related to stepping cadence of the metronome (0.8 Hz). To assess changes in M/L step regulation associated with increasing visual stimulus amplitude, average step width and step-width variability were measured and compared with changes in visual sensitivity. Both gait parameters were measured over the last 5 cycles of walking with the visual multi-sine stimulus including over 300 steps of walking. The first cycle was not included in the analysis to avoid capturing the transient response. Step width was measured at each step using heel marker coordinate positions as the absolute distance between the left and right heel markers at each heel strike of the trial. A heel strike event was determined as the point in time in which the heel marker was at the peak anterior position in the anterior/posterior direction. Average step width was calculated from the step width values over the entire trial. Step-width variability was calculated as the standard deviation of their step widths throughout the trials. Since participants performed 2 trials for each visual stimulus amplitude, each gait parameter was calculated as an average of measures from the two trials. Changes in each gait parameter and changes in visual sensitivity were calculated as differences between the 6 and 15 degree peak-to-peak conditions.

## Measuring Visual Sensitivity to Balance

To quantify the dynamic characteristics of the sensorimotor control process for balance during walking (visual sensitivity), we used methods previously developed to study standing balance (Peterka, 2002) and adapted them to be applicable for walking. Figure 3 (A) shows the 15-degree amplitude multi-sine stimulus in the time domain with a corresponding exemplary participant’s lateral CoM movement averaged over the last 5 stimulus cycles in Figure 3 (C). Both the stimulus and the lateral CoM movement are transformed from the time domain to frequency domain using a Fourier transform shown in Figure 3 (B) and (D), respectively. Red stem plots highlight the ten sinusoid frequencies that compose the multi-sine stimulus. Highlighted in Figure 3 (D) is the amplitude spectrum of the CoM movement resulting from participants following the metronome at 0.8 Hz (96 bpm). For the frequency response function analysis, the desired outcome is to analyze body motion caused by the visual stimulus regardless of the natural body sway that occurs in the medial/lateral direction due to the shifting of weight from one leg to another. The metronome drove the natural body sway well above the driving frequencies of the visual stimulus.

The frequency domain analysis consisted of calculating a frequency response function (FRF) for each participant at each stimulus amplitude. An FRF is a complex valued function that characterizes the stimulus/response behavior of the system and can be expressed as gain (sensitivity given by the ratio of response amplitude to stimulus amplitude) and phase (relative timing of the response) measures as a function of the 10 frequency components of the stimulus. The FRFs were calculated following the methods described in Pintelon & Schoukens (2012).

Specifically, the FRF 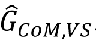 relating the M/L CoM displacement to the visual stimulus (VS) is given by:

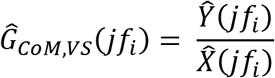

Where 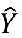 and 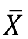 are the CoM and visual stimulus, respectively, cycle-averaged discrete Fourier transforms obtained by averaging across the last 5 cycles from the two trials (first cycle in each trial not included since it is a transient response) at the 10 stimulus frequencies, *fi*, and *j* is the imaginary number. The discrete Fourier transforms were calculated using the Matlab ‘*fft*’ function (Mathworks, Natick MA, Matlab version 2020a). The FRF gain (units of cm/deg) is given by the absolute value of 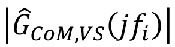 and the phase function (units deg) is given by:

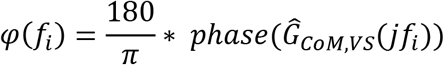

using the Matlab function ‘*phase*’ to calculate the arc tangent of the ratio of the imaginary to real components of 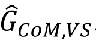

An overall measure of the sensitivity of balance to visual motion, *Sv*, was calculated by averaging the FRF gains across the frequencies corresponding to n = 10 stimulus frequency components:

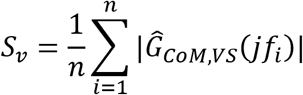

The stimulus amplitude-dependent change in sensitivity was calculated as the difference between *Sv* measured at the 6 and 15 degree visual stimulus amplitudes.

For each subject at each stimulus amplitude a coherence function was calculated whose values range from 0 to 1 at each stimulus frequency. Coherence values less than 1 indicate the presence of noise in measurements or inputs other than the visual stimulus contributing to the response. The coherence function, γ^2^, was calculated by:

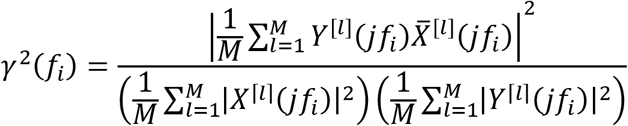

Where the discrete Fourier transforms of the stimulus *X* and CoM displacement response, *Y*, of each cycle were averaged over the *M* = 10 individual stimulus and response cycles, *l*, and 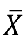 is the complex conjugate of 𝑋𝑋.

## Remnant Sway

To quantify the magnitude of body sway not driven by the visual multi-sine, we first calculated the scaled power spectrum of the average cycled M/L CoM trajectory (units of cm^2^/hz) for each participant. We then analyzed 9 even harmonics of the 40-s stimulus cycle, spanning 2/40 Hz to 18/40 Hz, which include frequencies not stimulated by the visual multi-sine. These 9 values were: 1) summed together, 2) multiplied by the fundamental frequency (0.025Hz), then 3) square-rooted for a final value in units cm.

## Statistical Analysis

To address our hypotheses relating to visual reweighting, a two-way repeated measures ANOVA was run to test for: 1) any significant main effects for changes in visual sensitivity caused by changing the stimulus amplitude, 2) main effects of group on visual sensitivity, 3) and any interaction effect between stimulus amplitude and group for differences in the degree of visual down-weighting between the two groups. Post-hoc analyses were performed to check for any significant differences within and between groups.

To analyze any changes in the spatiotemporal gait parameters, a two-way repeated measures ANOVA was run for each parameter to test for: 1) any significant main effects for changes in gait parameters caused by increasing the stimulus amplitude, 2) main effects of group on gait parameters, 3) and to check for any interaction effect between stimulus amplitude and group for differences in the degree of changes in each gait parameter between the two groups. Post-hoc analyses were performed to check for any significant differences within and between groups.

To test whether a relationship exists between a participants’ degree of visual down-weighting to the degree of each gait parameter, a Pearson’s correlation was run to measure the strength of the relationship between the difference of visual sensitivity and difference of each gait parameter for each participant taken at the 6-degree amplitude condition and the 15 degree amplitude condition in both groups. R^2^ values were obtained to reveal how much of the variance in the changes of each spatiotemporal parameter are explained by the change in visual re- weighting as measured by change in visual sensitivty.

## Results

A total of 60 subjects completed the experiment including 30 healthy young adults and 30 healthy older adults. We first present walking metrics (speed, step width, and step-width variability) followed by results on visual response sensitivity and finally relationships between walking metrics and visual response sensitivity. The focus is on characterizing changes related to visual stimulus amplitude and on differences between younger and older adults.

## Walking metrics

Table 2 shows the mean and standard deviation of walking speed, step width, and step-width variability and statistical results from the ANOVA.

**Table 2:**
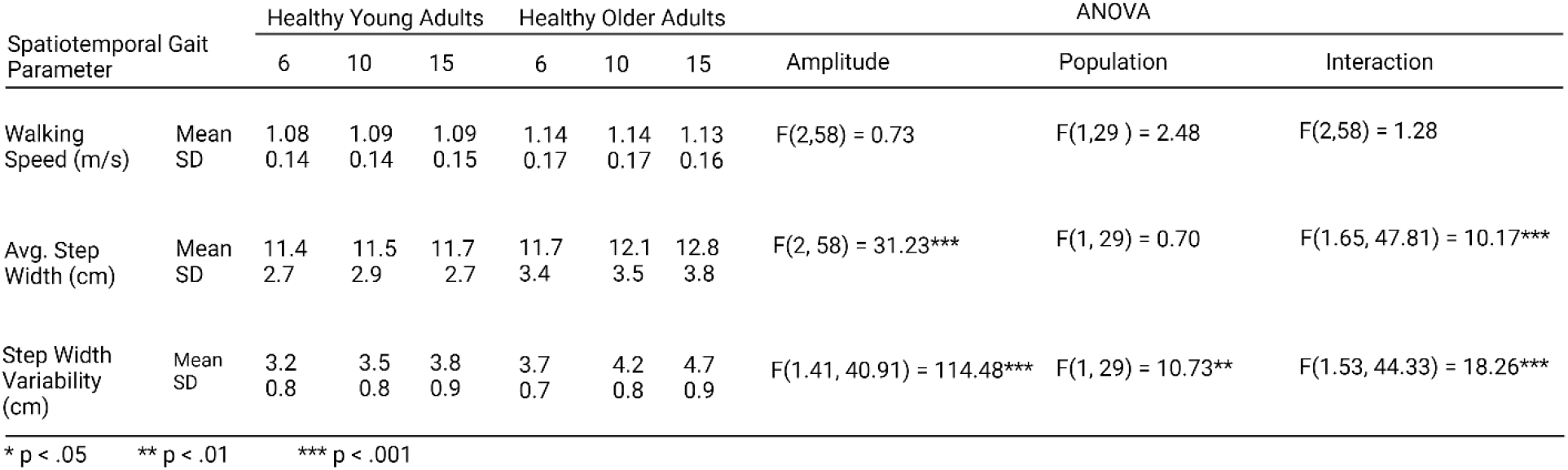
Spatiotemporal gait parameters and statistical results.

## Walking speed

Average walking speeds were not different between young and older adults (p = 0.126). Neither group showed variation in walking speed with visual stimulus amplitude and there was no interaction between the effects of visual stimulus amplitude and group indicating the visual stimulus did not affect walking speed in the two groups.

## Step Width

Step width generally increased with visual stimulus amplitude supported by a significant main effect of stimulus amplitude on average step width F(2,58) = 31.23, p < 0.001. Post-hoc pairwise t-tests with a Bonferroni correction revealed significant differences for young adults between stimulus amplitudes 6 and 15 and for all the amplitude conditions for older adults. There was no significant effect of group on step width. There was a significant interaction effect of visual stimulus amplitude by group on step width F(1.65,47.81) = 10.17; p < 0.001, indicating that changes in stimulus amplitude affected step width more for older adults.

## Step-Width Variability

Step with variability generally increased with increasing visual stimulus amplitude and was larger in older compared to younger adults, supported by a significant main effect of stimulus amplitude on step-width variability F(1.41,40.91) = 114.480, p < 0.001. Post-hoc pairwise t-tests with a Bonferroni correction revealed significant differences within groups for all the amplitude conditions with step-width variability increasing as stimulus amplitude increased. There was also a significant main effect of group on step-width variability F(1,29) = 10.73, p = 0.003. Post-hoc pairwise t-tests with a Bonferroni correction revealed significant differences between groups for all the amplitude conditions with step-width variability being significantly larger for older adults at each stimulus amplitude. There was a significant interaction between the effects of visual stimulus amplitude and group on step-width variability F(1.53,44.33) = 18.26; p < 0.001, indicating that changes in stimulus amplitude affected step-width variability more for older adults.

## Visual Responses

### Visual Evoked Sway

Figure 4 shows M/L CoM displacement averaged across stimulus cycles as a function of increasing amplitude of the visual multi-sine stimulus for both healthy young and older adults. Qualitatively, the time course of the CoM displacement resembled the time course of the visual stimulus (see Figure 3A) but generally increased with increasing stimulus amplitude (Figure 4). CoM movement was quantified by calculating the root mean square (RMS) value of the M/L CoM trajectory with results shown in Figure 5. The mean RMS value of CoM trajectories for young adults were 2.7 cm, 3.4 cm, and 4.3 cm during the 6, 10, and 15 degree conditions, respectively, and for older adults were 3.4 cm, 4.7 cm, and 6.0 cm, respectively.

**Figure 4:**
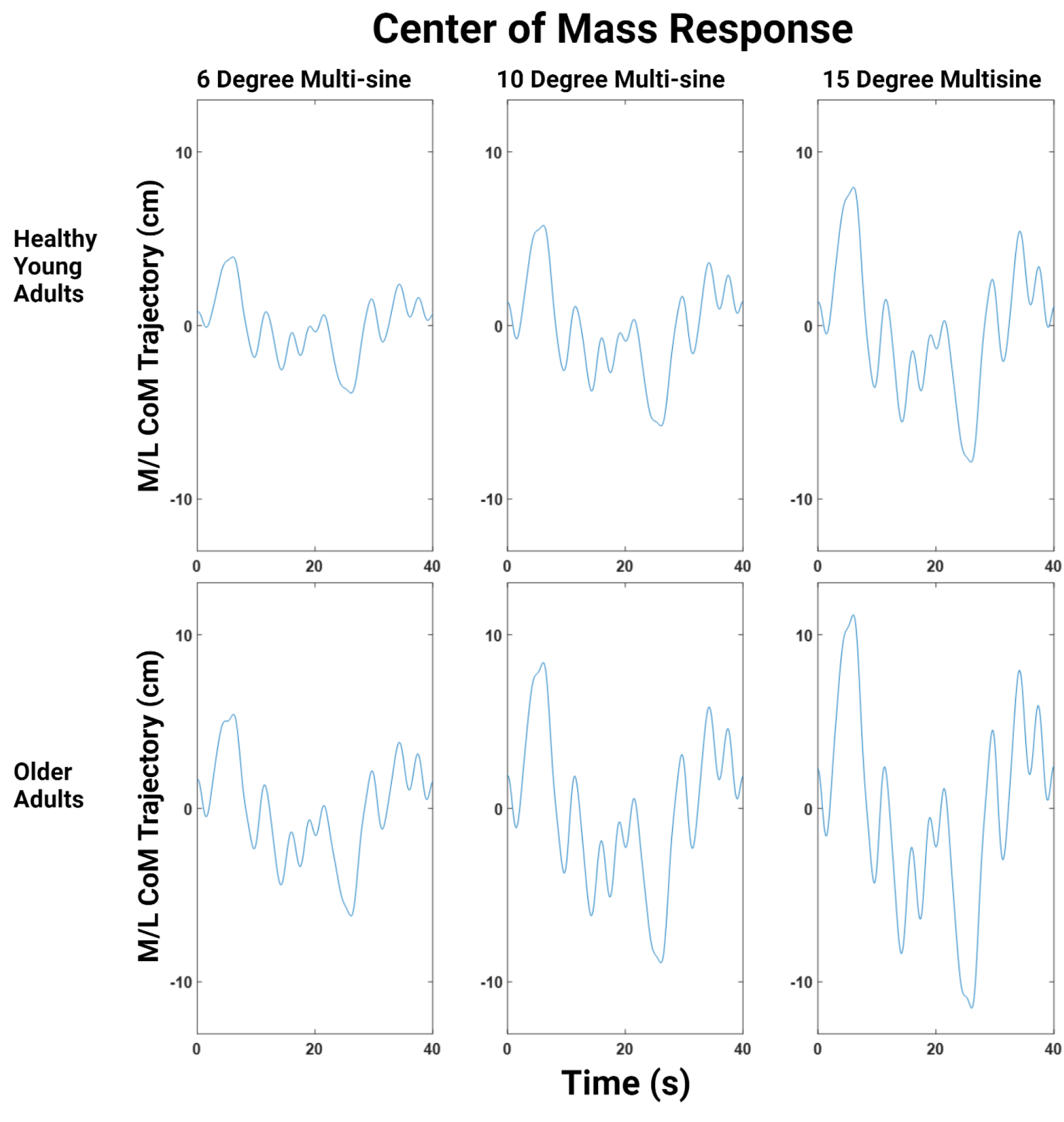
Average M/L CoM response as a function of increasing visual multi-sine stimulus amplitude. Top row - young adults; Bottom row - older adults.

**Figure 5:**
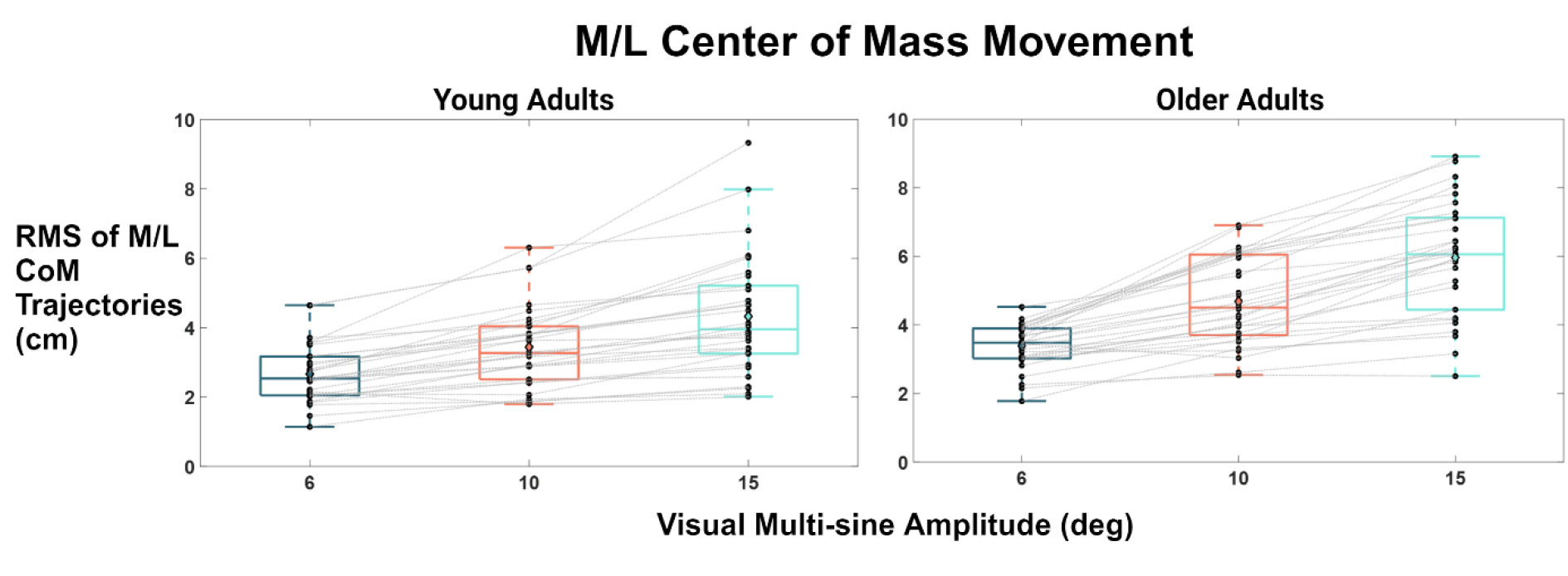
Root Mean Square (RMS) measures (cm) of the M/L CoM trajectory of body sway while walking and viewing the visual stimulus at the three stimulus amplitudes for young and older adults.

### Visual Sensitivity

Figures 6 and 7 show the FRFs relating M/L CoM movement to the visual stimulus. FRF gain measures were averaged across the 10 multi-sine frequencies to calculate visual sensitivity (*Sv*, cm/deg) to balance for each visual stimulus amplitude. Figure 8 shows the measures of visual sensitivity (*Sv*, cm/deg) for balance control for each stimulus amplitude comparing healthy young (A) and older adults (B). Visual sensitivity generally decreased as the stimulus amplitude increases for both young and older adults.

**Figure 6:**
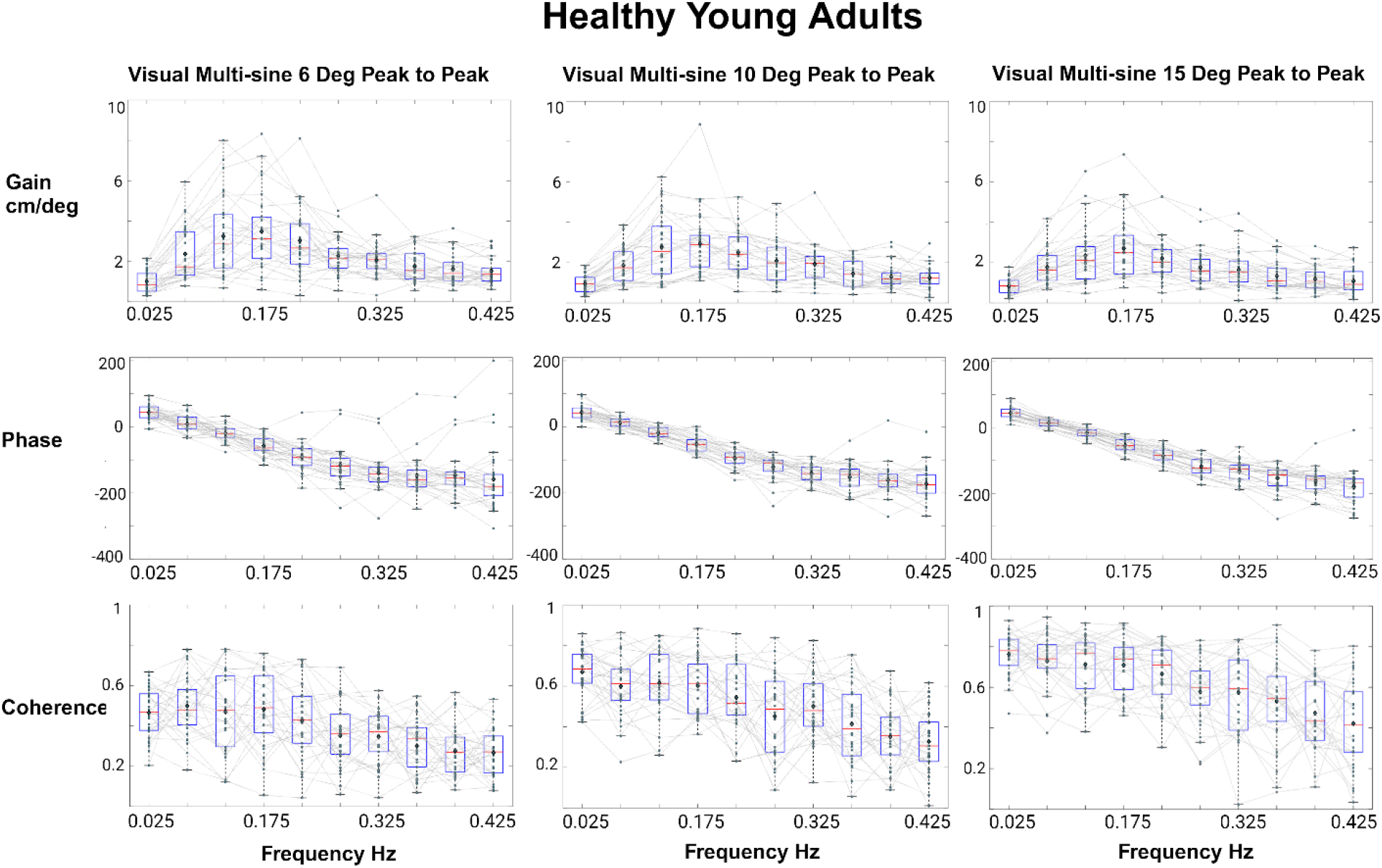
Shows the gain, phase, and coherence of the frequency response functions for healthy young adults measured over the 10 driving frequencies of the visual multi-sine stimulus at the three different multi-sine amplitudes: 6, 10, and 15 degrees peak-to-peak.

**Figure 7:**
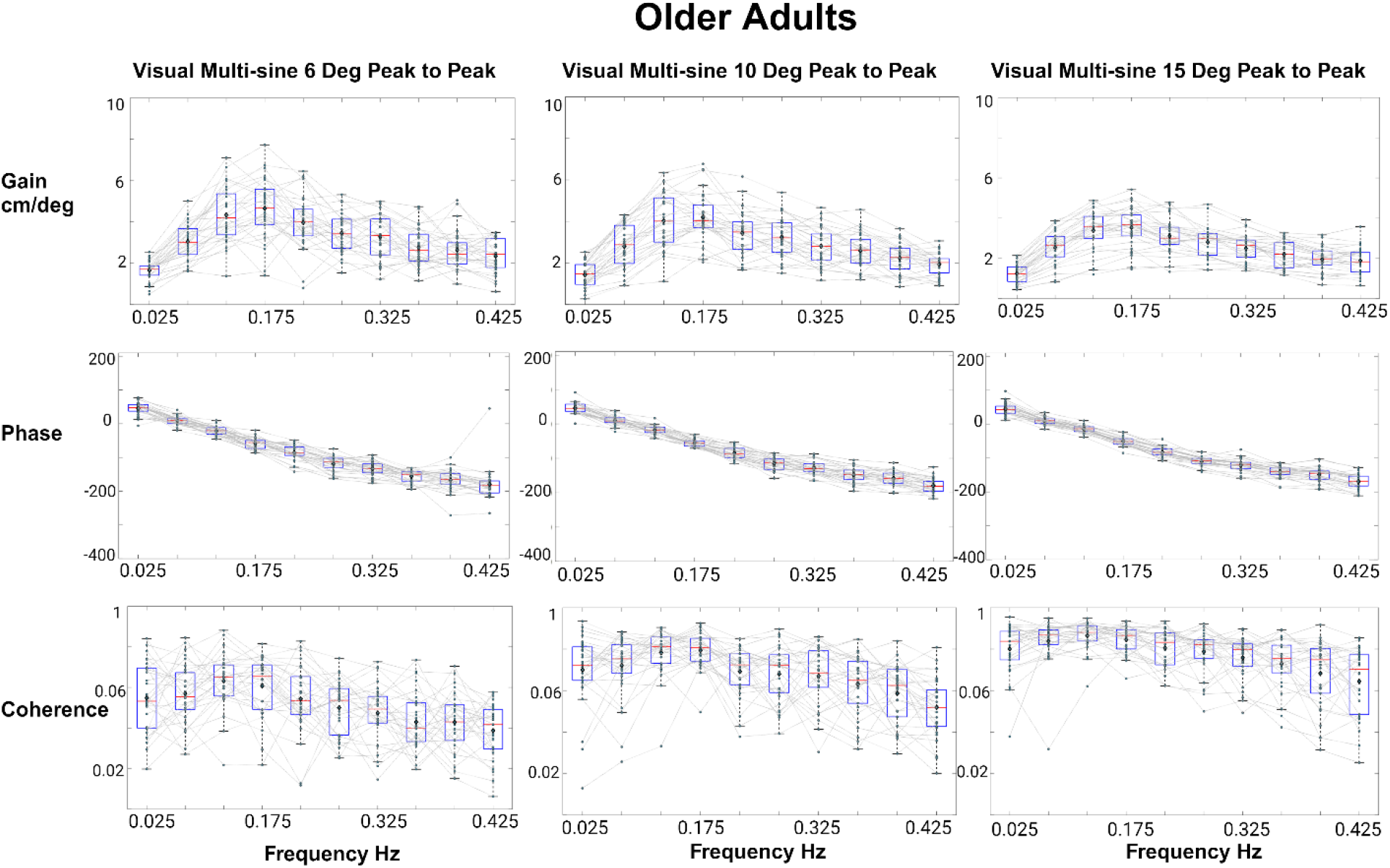
Shows the gain, phase, and coherence of the frequency response functions for healthy older adults measured over the 10 driving frequencies of the visual multi-sine stimulus at the three different multi-sine amplitudes: 6, 10, and 15 degrees peak-to-peak.

**Figure 8:**
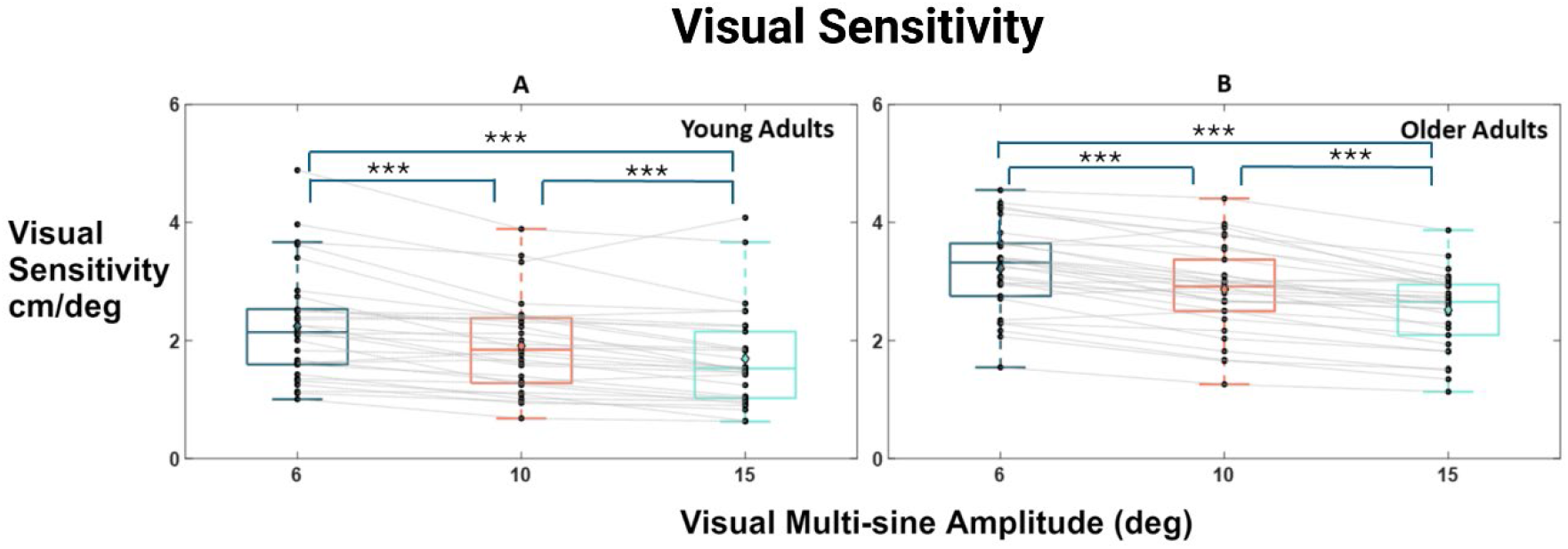
Compares the measures of visual sensitivity to M/L balance control as a function of increasing stimulus amplitude for both healthy young (A) and older adults (B). The *** symbols represent p values less than 0.001.

**Figure 9:**
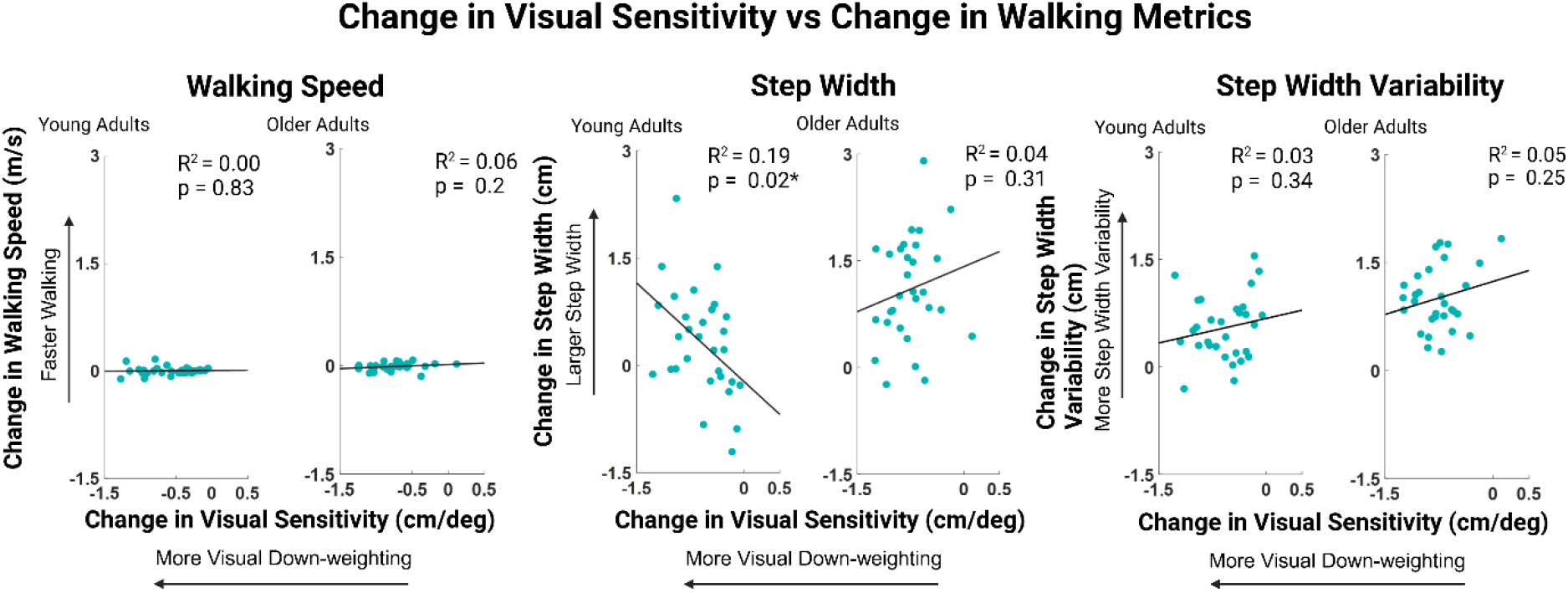
Scatter plots representing the relationship between the change in visual sensitivity to the change in walking metrics from the lowest stimulus amplitude (6 deg) to the largest (15 deg). Walking Metrics from left to right include walking speed (m/s), step width (cm), and step-width variability (cm). Pearson’s R2 and corresponding p values are listed above each correlation plot.

Simple main effects from the two-way repeated measures ANOVA showed that amplitude had a significant effect on visual sensitivity F(2,58) = 125.61, p < 0.001. Post-hoc pairwise t-tests with a Bonferroni correction revealed significant differences within groups for all the amplitude conditions, where visual sensitivity decreased as stimulus amplitude increased for each condition (p < 0.001). Simple main effects of the two-way repeated measures ANOVA showed a significant effect for group on visual sensitivity F(1,29) = 23.04, p < 0.001. Post-hoc analysis revealed significant differences between groups at each stimulus amplitude, where older adults had higher visual sensitivity at each amplitude (p < 0.001). Lastly, a two-way ANOVA revealed there was no significant interaction effect between the visual stimulus amplitude and group on visual sensitivity measures F(2,58) = 2.97; p < 0.06, revealing that changes in stimulus amplitude effected both groups similarly.

### Remnant sway

The remnant sway measures provided an indication of the variability of M/L CoM motion that was not accounted for by the stimulus-evoked CoM motion. Remnant sway values at the three stimulus amplitudes for both young and older adults are shown in Table 3. Remnant sway was slightly higher in older compared to younger adults (average 0.16 cm larger) although this result was not statistically significant. Neither group showed variation in remnant sway with visual stimulus amplitude and there was no interaction between the effects of visual stimulus amplitude and group indicating the visual stimulus affected remnant sway similarly between the two groups (p = 0.34).

**Table 3:** Includes average power of medial/lateral body sway found at the even harmonics of the visual multi-sine stimulus including 9 frequencies ranging from 0.05Hz - 0.45Hz.

| Stimulus Evoked Sway |  | Healthy Young Adults |  |  | Healthy Older Adults |  |  | ANOVA |  |  |
| --- | --- | --- | --- | --- | --- | --- | --- | --- | --- | --- |
|  |  | 6 | 10 | 15 | 6 | 10 | 15 | Amplitude | Population | Interaction |
| Visual Sensitivity (cm/deg) | Mean | 2.31 | 1.93 | 1.72 | 3.29 | 2.89 | 2.54 | F(2,58) = 125.61 *** | F(1,29) = 23.04 *** | F(2,58) = 2.97 |
|  | SD | 0.95 | 0.78 | 0.81 | 0.75 | 0.76 | 0.65 |  |  |  |
| Remnant Sway (cm) | Mean | 2.07 | 2.06 | 2.09 | 2.24 | 2.19 | 2.26 | F(2,58) = 0.60 | F(1,29) = 0.942 | F(2,58) = 0.09 |
|  | SD | 0.75 | 0.72 | 0.78 | 0.46 | 0.49 | 0.55 |  |  |  |
| * p < .05 | ** p < .01 | *** p < .001 |  |  |  |  |  |  |  |  |

### Relationships between Visual Sensitivity Changes and Changes in Walking Metrics

Pearson’s correlations for both young and older adults were used to compare the stength of the relationship between the visual stimulus amplitude-dependent change in visual sensitivity and change in walking speed, average step width, and step-width variability (Figure 9). Change was defined as the value of the metric for the 15 deg stimulus minus the value for the 6 deg stimulus. No significant correlations were found between changes in visual sensitivity and changes in walking speed for both young adults (R^2^ = 0.00; p = 0.83) and older adults (R^2^ = 0.06; p = 0.20). A weak, but significant negative correlation was found between changes in visual sensitivity and changes in average step width for young adults (R^2^ = 0.19; p = 0.02) indicating the individuals that showed a larger change in visual sensitivity (i.e., larger negative values) with increasing stimulus amplitude tended to have a larger positive change (increase) in step width.

For older adults the correlation had the opposite sign (positive correlation) but was not significant (R^2^ = 0.04; p = 0.31). Correlations were positive between changes in visual sensitivity and changes in step-width variability for both young adults (R^2^ = 0.03; p = 0.34) and older adults (R^2^ = 0.05; p = 0.25), but neither was significant.

## Discussion

Our study investigated the change in emphasis of vision for maintaining M/L balance while walking under varying visual stimulus conditions. Healthy young and older participants walked along a self-paced treadmill while viewing a virtual environment that tilted in the frontal plane using a multi-sine stimulus at three different maximum amplitudes: 6, 10, and 15 degrees peak-to-peak. Frequency response functions comparing the visual stimulus to the M/L CoM movement showed that for both populations, there was a decrease in visual sensitivity for balance control as the visual stimulus increased in amplitude, suggesting that neurotypical adults decreased reliance on vision for walking balance control as a visual stimulus increased in magnitude. However, this decrease in visual sensitivity was not accompanied by an increase in remnant body sway as hypothesized. Average step width and step-width variability significantly increased as a function of increasing visual stimulus amplitude for both groups, while waking speed was unaffected.

Older adults showed significantly higher visual sensitivity at all three amplitude conditions compared to young adults, suggesting that older adults rely more on vision for balance control compared to young adults. This finding is consistent with current literature reviewed by Osoba et al (2019). Interestingly, although visual sensitivity to balance was overall higher in older adults, older adults commonly stated that they did not perceive any changes in the visual environment during the multi-sine stimulus conditions, particularly the 6-degree amplitude condition. Young adults did not state any lack of perceptual awareness that the screen was tilting. Visual motion detection thresholds have been shown to be higher in older adults (Conlon et al, 2017; Snowden and Kavanagh, 2006), which may explain the lack of perceptual awareness of the moving scene. But despite this, older adults consistently demonstrated greater responses to the visual stimuli.

Our hypothesis that the amplitude-dependent changes in visual sensitivity was attributable to sensory reweighting is by analogy to previous results from experiments investigating the control of balance during stance that have shown similar amplitude-dependent sensitivity changes in response to tilts of the visual scene and the stance surface (Peterka, 2002; Peterka et al., 2018). In these previous studies the sensory weights were estimated by adjusting parameters of a mathematical feedback control model of the balance system to optimally account for experimental data. An assumption of the model was that the sensory weights for all sensory systems contributing to balance sum to one meaning that an amplitude-dependent decrease in the sensory weight from one sensory system implied that the sensory weights of other sensory systems increase. This was confirmed in a study using surface-tilt stimuli combined with galvanic stimuli that evoked M/L sway to investigate changes in proprioceptive and vestibular contributions to balance as a function of surface tilt amplitude (Cenciarini and Peterka, 2006) and also in a study by Hwang et al. (2014) that used combinations of visual motion, Achillies tendon vibration, and galvanic vestibular stimulation to investigate intramodal and intermodal effects on response sensitivity. Similarly, experimental manipulations that altered the availability of sensory orientation information from vision (lights on or off) or proprioception (fixed or sway-referenced stance surface) evoked changes in response sensitivity to M/L balance perturbations consistent with sensory reweighting (Assländer and Peterka, 2016).

The Assländer and Peterka study (2016) and another study investigating proprioceptive/vestibular reweighting (van der Kooij and Peterka, 2011) showed that sway variability and remnant sway, respectively, increased with manipulations that evoked sensory reweighting. That is, increased sway variability is a consequence of the re-weighting shift toward utilization of less precise (i.e., noisier) sensory sources to compensate for the disruption caused by a balance perturbation applied to one sensory system. Therefore, we hypothesized that we would see an increase in remnant sway with increasing stimulus amplitude in our walking experiments, but this was not found (Table 3).

Here we discuss two possible reasons why remnant sway did not show the expected increase with stimulus amplitude. First, it could be that the amplitude-dependent decrease in visual sensitivity was not caused by down-weighting of visual sensory contributions to balance but by factors affecting the motor actions that respond to balance perturbations. In standing balance control, motor activation factors (i.e., factors that determine the magnitude of corrective torque generation) have been shown to have an influence on sensitivity to balance disturbances (Campbell et al., 2022). Specifically, a subject who generates a greater amount of corrective motor action can appear to be mechanically ‘stiffer’ than a subject with lower corrective action with the result that a given balance perturbation will produce less sway in the stiffer subject. If, on average, stiffness increased with increasing stimulus amplitude then visual sensitivity would decrease without there necessarily being a change in sensory weighting. Future experiments that simultaneously present two stimuli to perturb different sensory systems could be used to test for explicit reweighting effects where an increase in response sensitivity to one sensory disturbance is accompanied by a decrease in response sensitivity to the other sensory disturbance.

A second reason for the absence of an amplitude-dependent effect on remnant sway may be related to the source of the sway variability. In standing balance control there is evidence that sensory noise, as opposed to motor noise, is the dominant contributor to sway variability (van der Kooij and Peterka, 2011). Therefore, the remnant sway is very sensitive to changes in sensory weights when the balance system shifts toward reliance on less precise (nosier) sensory sources. However, if the source of sway variability during walking is dominated by motor variability rather than sensory variability, then little or no detectable change in sway variability would be expected even if sensory reweighting contributed to changes in response sensitivity. The understanding of the sensory versus motor noise contributions to sway variability during stance relied on the application of system identification and modeling methods. In principle, these methods could be applied to balance during walking. This would require having models that represent the much more complex aspects of balance control during walking that take into consideration the sensory contributions, which may vary across the gait cycle (Bent et al., 2004, Duysens et al., 1995, Logan et al., 2010), the multiple motor mechanisms that contribute to corrective actions (Reimann et al., 2018), and the influence of gait parameters (e.g., step width, walking speed, step timing).

We calculated and averaged the filtered CoM trajectories (CoM Response) during each visual stimulus amplitude condition and found that although the sensitivity to the visual stimulus decreased relative to the stimulus amplitude, the overall movement caused by the virtual perturbations increased with increasing stimulus amplitude (Figures 4 & 5). This increased response to the visual stimulus was accompanied by an increase in coherence observed over the three amplitudes (Figures 6 & 7). Increased coherence represents an increase in the signal-to-noise ratio between the visual stimulus and the M/L CoM movement.

Increased CoM movement was accompanied by increased average step width and step-width variability (Table 2), supporting previous work that destabilizing environments via visual and support surface perturbations cause increased step-width variability in healthy young and older adults (McAndrew et al. 2010; Franz et al., 2015; Kazanski et al., 2020). According to the predictive model set forth by Wang and Srinivasan (2014), the CoM position at midstance predicts the variance (∼80%) of the following step placement, so a sensory perturbation that adds noise to estimating vertical alignment would inherently add variability to the CoM trajectory and thus add variability to step placement. We were interested, however, in exploring whether any relationship existed between the degree of visual sensitivity changes and resulting changes in gait parameters, and to observe whether visual sensitivity changes, and their potential association with sensory reweighting, act as a neural mechanism that contributes to M/L stability via step regulation. Correlation coefficients between the change in visual sensitivity and the change in average step width showed a significant but weak negative correlation for young adults while change in visual sensitivity and step-width variability showed non-significant positive correlations for both groups (Figure 9). Although we had no a priori hypothesis, our thought was that individuals whose balance control system was better at reducing the influence of the visual disturbance would also show lower destabilizing effects of the increasing visual perturbation.

Overall, for younger subjects, a greater ability to change (reduce) visual sensitivity with increasing stimulus amplitude was associated with a greater change in step width (toward wider step widths which may contribute to stability) and with a smaller change in step-width variability. Although not significant, the smaller change in step-width variability possibly hints at the effectiveness of the change in visual sensitivity, or its association with step width changes, in reducing instability at larger visual stimulus amplitudes. However, older adults did not show a significant change in step width or step-width variability with change in visual sensitivity.

Identifying normative data for sensory sensitivity for balance control can be a potential tool for understanding how an individuals’ nervous system utilizes sensory information for balance control. For example, here we probed visual processing and found that young individuals moved 2.31 cm per degree of visual tilt. An individual displaying significantly higher or lower sensitivities to visual tilts may indicate the presence of sensory or central nervous system disorders. For example, individuals with cerebral palsy are more perturbed by visual tilts compared to young adults, and this behavior is associated with somatosensory deficits (Sansare et al. 2022). Establishing normative data for sensory sensitivity for balance control in diverse populations (including those with peripheral sensory deficits and central neurological disorders) can provide insight into how individuals process sensory information and compensate for environmental changes. This insight could inform the development of future rehabilitation strategies.

## Conclusion

Our study found that healthy young and older adults become less sensitive to visual perturbations as the amplitude of the visual stimulus increases. This decrease in sensitivity potentially is analogous to the sensory reweighting hypothesis observed in standing balance control as a neural mechanism for upright M/L balance control during walking although future effort is needed to explore the influence of other motor control and biomechanical factors that likely influence the sensitivity to balance perturbations. We provide a measure of visual sensitivity that could be adapted to probe other systems such as the vestibular system via galvanic vestibular stimulation. Given that most falls occur during walking, the risk of falling may be informed by better understanding the sensitivity of an individual to disturbances in a sensory modality and the factors that influence this sensitivity.

